# Regulation of melanocyte development by ligand-dependent BMP signaling underlies oncogenic BMP signaling in melanoma

**DOI:** 10.1101/708610

**Authors:** Alec K. Gramann, Arvind M. Venkatesan, Melissa Guerin, Craig J. Ceol

**Affiliations:** Program in Molecular Medicine and Department of Molecular, Cell, and Cancer Biology, University of Massachusetts Medical School, 368 Plantation St, Worcester, MA 01605, USA

**Keywords:** Melanoma, melanocyte, BMP signaling, neural crest, specification, mitf, mitfa, zebrafish

## Abstract

Preventing terminal differentiation is important in the development and progression of many cancers including melanoma. Recent identification of the BMP ligand *GDF6* as a novel melanoma oncogene showed *GDF6*-activated BMP signaling suppresses differentiation of melanoma cells. Previous studies have identified roles for *GDF6* orthologs during early embryonic and neural crest development, but have not identified direct regulation of melanocyte development by GDF6. Here, we investigate the BMP ligand *gdf6a*, a zebrafish ortholog of human *GDF6*, during the development of melanocytes from the neural crest. We establish that the loss of *gdf6a* or inhibition of BMP signaling during neural crest development disrupts normal pigment cell development, leading to an increase in the number of melanocytes and a corresponding decrease in iridophores, another neural crest-derived pigment cell type in zebrafish. This shift occurs as pigment cells arise from the neural crest and depends on *mitfa*, an ortholog of *MITF*, a key regulator of melanocyte development that is also targeted by oncogenic BMP signaling. Together, these results indicate that the oncogenic role ligand-dependent BMP signaling plays in suppressing differentiation in melanoma is a reiteration of its physiological roles during melanocyte development.

## Introduction

Tumor differentiation status is often an important prognostic factor in cancer. For many cancer types, tumors that are less differentiated are associated with a higher grade and worse prognosis compared to more differentiated tumors, which often follow indolent courses (Hoek et al., 2006; Rosai & Ackerman, 1979). In order to adopt a less differentiated state, a common event in cancer is downregulation of factors that drive differentiation of adult tissues (Chaffer et al., 2011; Dravis et al., 2018). This loss of pro-differentiation factors is often coupled with an upregulation of other factors that are associated with embryonic or progenitor states (Caramel et al., 2013; Tulchinsky, Pringle, Caramel, & Ansieau, 2014). Thus, many de-differentiated and high-grade cancers have gene expression profiles associated with early development (O’Brien-Ball & Biddle, 2017).

Developmental factors and pathways co-opted by cancers are often related to vital cellular functions, such as proliferation, migration, and differentiation (Caramel et al., 2013; Casas et al., 2011; McConnell et al., 2019; Perego et al., 2018). Furthermore, the embryonic origin of specific tissues can impact the aggressive phenotypes tumors are able to acquire (Carreira et al., 2006; Gupta et al., 2005; Hoek & Goding, 2010). In the case of melanoma, the cell of origin, the melanocyte, is derived from the neural crest, a highly migratory population of embryonic cells. Thus, melanomas are prone to early and aggressive metastasis, associated with the expression of neural crest migratory factors (Liu, Fukunaga-Kalabis, Li, & Herlyn, 2014). Additionally, melanomas lacking differentiation exhibit more aggressive characteristics and are broadly more resistant to therapy (Fallahi-Sichani et al., 2017; Knappe et al., 2016; Landsberg et al., 2012; Mehta et al., 2018; Muller et al., 2014; Shaffer et al., 2017; Zuo et al., 2018). While differentiation status is evidently important in the course of disease, the mechanisms by which melanomas and other cancers remain less differentiated are poorly understood. Since many of the factors associated with a lack of differentiation in these cancers are expressed and apparently function during embryogenesis, elucidating the developmental roles of these factors can give insight into their behaviors and roles in tumorigenesis and progression.

A key pathway involved in early development and development of the neural crest is the bone-morphogenetic protein (BMP) pathway (reviewed in Kishigami & Mishina, 2005). The BMP pathway is activated by BMP-ligands binding to BMP receptors, which can then phosphorylate SMAD1, SMAD5, and SMAD8 (also called SMAD9). Phosphorylated SMAD1/5/8 associates with co-SMAD4, forming a complex that can translocate to the nucleus and regulate expression of target genes. BMP signaling is important in early embryonic dorsoventral patterning and induction of the neural crest (Garnett, Square, & Medeiros, 2012; Hashiguchi & Mullins, 2013; McMahon et al., 1998; Schumacher, Hashiguchi, Nguyen, & Mullins, 2011). Following neural crest induction, BMP signaling has been implicated in patterning within the neural crest and surrounding tissues, as well as development of nervous system- and musculoskeletal-related neural crest lineages (Hayano, Komatsu, Pan, & Mishina, 2015; McMahon et al., 1998; Nikaido, Tada, Saji, & Ueno, 1997; Reichert, Randall, & Hill, 2013; Valdivia et al., 2016). While many developmental functions of BMP signaling are well characterized, the relationship of BMP signaling to the development of pigment cells from the neural crest is poorly understood.

Our laboratory recently identified a BMP ligand, *GDF6,* that acts to suppress differentiation and cell death in melanoma (Venkatesan et al., 2018). We found that *GDF6*-activated BMP signaling in melanoma cells represses expression of *MITF*, a key regulator of melanocyte differentiation, leading to a less differentiated state. Here, we investigate the role of *GDF6* and the BMP pathway in development of pigment cells in zebrafish. We show that BMP signaling regulates fate specification of neural crest-derived pigment cell lineages and suppresses expression of *mitfa*, an ortholog of *MITF*. Furthermore, we show that disrupting BMP signaling alters fate specification between melanocyte and iridophore populations in the zebrafish. We determine that this shift in fate occurs at the level of an *mitfa-*positive pigment progenitor cell, and that BMP signaling acts through *mitfa* to direct *mitfa*-positive pigment progenitor cells to a specific fate. Altogether, these findings suggest pathologic BMP signaling in melanoma is a reiteration of normal physiologic function of BMP signaling during melanocyte development.

## Results

### Loss of *gdf6a* leads to an increase in adult pigmentation

To understand potential functions of *gdf6a* in the melanocyte lineage, we first determined if any alterations in pigment pattern were present in animals lacking *gdf6a*. In these studies, we used the *gdf6a*^s327^ allele, hereafter referred to as *gdf6a(lf),* which encodes an early stop codon and has previously been shown to cause a complete loss of *gdf6a* function (Gosse & Baier, 2009). Previous studies have identified early roles for *gdf6a* during initial embryonic patterning, including dorsoventral patterning immediately following fertilization, thus *gdf6a(lf)* mutants have significantly decreased viability during the first 5 days post fertilization (Sidi, Goutel, Peyrieras, & Rosa, 2003). However, we found that a small proportion of *gdf6a(lf)* animals are able to survive early development and progress to adulthood. These *gdf6a(lf)* adult zebrafish had increased pigmentation when compared to wild-type zebrafish (Figure 1A). Furthermore, *gdf6a(lf)* adult zebrafish had qualitative disruption of the normal pigment pattern of both stripe and scale-associated melanocytes, and a significant increase in the number of scale-associated melanocytes as well as the overall scale area covered by melanin (Figure 1A,1B). These results indicate that *gdf6a(lf)* mutants have melanocyte defects.

**Figure 1.**
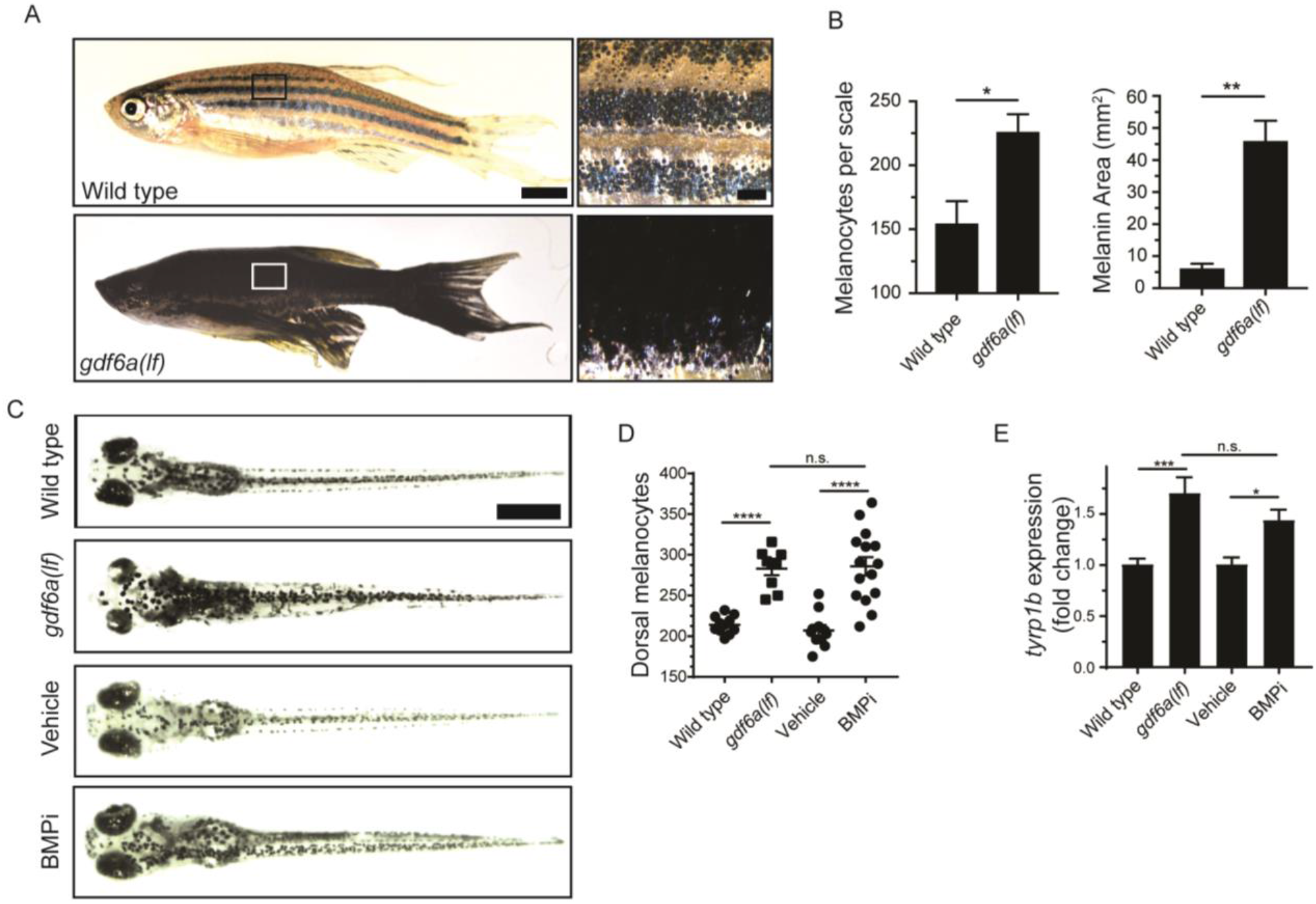
*gdf6a* loss or BMP inhibition causes the development of supernumerary melanocytes. (A) Images of wild-type and *gdf6a(lf)* adult zebrafish, scale bar = 4 mm, inset scale bar = 1 mm. (B) Quantification of number of melanocytes (left) and scale pigmentation using melanin coverage (right), n = 3 scales per group. (C) Wild-type and *gdf6a(lf)* embryos imaged at 5 days post fertilization (DPF); vehicle- and BMPi-treated embryos imaged at 5 DPF. Scale bar = 1 mm. Animals were treated with epinephrine prior to imaging. (D) Quantification of dorsal melanocytes per animal in 5 DPF wild-type, *gdf6a(lf)* mutant, vehicle-, and BMPi-treated embryos. n = 11, 9, 11, and 15, respectively. (E) Expression of *tyrp1b* by qPCR in wild-type, *gdf6a(lf)* mutant, vehicle-, and BMPi-treated embryos. n = 5-6 for each group. Error bars represent mean +/-SEM. P-values were calculated using Student’s t-test in panel B and one-way ANOVA with Tukey’s multiple comparisons test in panels D and E, * P<0.05, ** P<0.01, *** P<0.001, **** P<0.0001, n.s., not significant.

### Loss of *gdf6a* or inhibition of BMP signaling leads to an increase in embryonic melanocytes

Since zebrafish develop their adult pigment pattern during metamorphosis, it is possible *gdf6a* acts during this stage to change adult pigmentation, and not during initial pigment cell development in embryogenesis (D. M. Parichy & Spiewak, 2015; Patterson & Parichy, 2013; Quigley et al., 2004). To address this issue, we investigated whether *gdf6a(lf)* caused embryonic pigmentation changes and, if so, whether any such changes were BMP-dependent. We crossed *gdf6a(lf)* heterozygotes and, in randomly selected progeny, quantified the number of melanocytes that developed by 5 days post-fertilization (DPF). Following melanocyte quantification, we determined the genotype of each embryo. In parallel, we treated wild-type zebrafish during the period of neural crest induction and melanocyte specification (12 to 24 hours post fertilization) with a small molecule BMP inhibitor, DMH1, hereafter referred to as BMPi, and performed the same quantification of embryonic melanocytes (Hao et al., 2010). *gdf6a(lf)* homozygous animals developed approximately 40% more dorsal melanocytes by 5 DPF, when compared to sibling wild-type animals and *gdf6a(lf)* heterozygotes (Figures 1C,1D and S1A). *gdf6a(lf)* animals also showed increased expression of *tyrp1b*, a marker of differentiated melanocytes, which is consistent with an increase in melanocyte number (Figure 1E). Furthermore, treatment with BMPi phenocopied the melanocyte changes observed in *gdf6a(lf)* mutants, coupled with a similar increase in expression of *tyrp1b* (Figures 1D and 1E). We observed a similar increase in total body melanocytes, indicating that there is an overall increase in melanocyte development instead of a failure of migration leading to a specific increase in dorsal melanocytes (Figure S1B). These results indicate *gdf6a*-activated BMP signaling normally acts in embryos to limit melanocyte development.

### *gdf6* ortholog expression during neural crest development

Numerous BMP ligands are expressed during early embryogenesis and participate in multiple facets of development, including neural crest induction. It was previously shown that multiple BMP ligands are activated during zebrafish neural crest development (Reichert et al., 2013). Of those ligands investigated, only *gdf6a* and *bmp6* were expressed in the neural crest, and only *gdf6a* activated BMP signaling within neural crest cells. An additional study identified dorsal expression of a zebrafish paralog of *gdf6a*, *gdf6b,* indicating it could potentially act in the neural crest (Bruneau & Rosa, 1997). We verified *gdf6b* expression is restricted to the neural tube, and further determined *gdf6b* loss of function has no impact on pigment cell development by generating a *gdf6b* mutant, hereafter referred to as *gdf6b(lf)*, and counting embryonic melanocytes (Figure S1D, S1E, S1F and S1G). We generated double mutants for both *gdf6a(lf)* and *gdf6b(lf)* to assess whether these paralogs functioned redundantly or could compensate for the loss of one another. Unfortunately, *gdf6a(lf); gdf6b(lf)* double mutants had significant morphologic defects and decreased viability such that we could not adequately compare melanocyte numbers in these animals (Figure S1H and S1I). However, because there were no pigmentation defects in *gdf6b(lf)* mutants and *gdf6a(lf)* pigmentation defects were the same severity as observed in animals treated with a pan-BMP inhibitor, it is likely that most, if not all, effects of BMP signaling on melanocyte development are directed by *gdf6a*.

### BMP inhibition increases *mitfa-*positive pigment cell progenitors in the neural crest

We sought to determine the mechanism by which BMP signaling inhibits melanocyte development in embryos. Based on our experiments using BMPi, we suspected BMP signaling acts during pigment cell development from the neural crest to prevent an increase in melanocytes. Following induction, neural crest cells undergo proliferation, followed by fate restriction and specification, in which individual cells become less and less multipotent until a single possible fate remains (Jin, Erickson, Takada, & Burrus, 2001; Lewis et al., 2004; Nagao et al., 2018). In many cases, specification to the ultimate lineage is determined by activation of an individual or a group of lineage-specific factors (Sauka-Spengler, Meulemans, Jones, & Bronner-Fraser, 2007). For pigment cells, fate specification is dependent on integration of many signaling factors, including BMP and Wnt signaling, as well as key transcription factors, such as *SOX*-, *PAX*-, and *FOX*-family transcription factors (Garnett et al., 2012; Ignatius, Moose, El-Hodiri, & Henion, 2008; Lister et al., 2006; Sato, 2005; Southard-Smith, Kos, & Pavan, 1998; Thomas & Erickson, 2009). In zebrafish, specification of the pigment cell lineage depends on upregulation of *sox10* and downregulation of factors inhibiting differentiation, such *as foxd3* (Curran et al., 2010; Curran, Raible, & Lister, 2009; Dutton et al., 2001). Following *sox10* upregulation, a subset of *sox10*-positive cells can activate pigment lineage markers associated with melanocytes, iridophores, and xanthophores (Elworthy, Lister, Carney, Raible, & Kelsh, 2003; Fadeev, Krauss, Singh, & Nusslein-Volhard, 2016; Nagao et al., 2018; Nord, Dennhag, Muck, & von Hofsten, 2016; Petratou et al., 2018). *mitfa* is a key factor that is expressed early in pigment progenitor cells (Lister, Robertson, Lepage, Johnson, & Raible, 1999). Based on this framework, we hypothesized two potential mechanisms by which supernumerary melanocytes are generated: 1) an increase in proliferation of either neural crest cells or pigment progenitor cells, or 2) an increase in the proportion of neural crest cells that are specified to become pigment progenitor cells. To assess changes in proliferation of neural crest cells and pigment cells, we analyzed cell cycle profiles using flow cytometry. Embryos expressing reporters for neural crest cells (*Tg(crestin:eGFP))* or pigment progenitor cells *(Tg(mitfa:eGFP))* were treated with BMPi from 12 to 24 HPF, during neural crest development and specification (Curran et al., 2009; Kaufman et al., 2016). Embryos were dissociated, stained with DAPI, and analyzed for DNA content of neural crest cells or pigment progenitor cells as defined by the fluorescent GFP marker (Figure S2A). We observed no increase in the percent of S/G2/M cells in either population, indicating no apparent change to proliferation in either neural crest cells or pigment progenitor cells (Figure S2B, S2C). Without an obvious increase in proliferation, we tested the hypothesis that a change in specification results in increased melanocytes. To assess changes in specification of neural crest cells into pigment progenitor cells, we utilized reporter embryos marking neural crest cells in red (*Tg(crestin:mCherry))* and pigment progenitor cells in green (*Tg(mitfa:eGFP*)) (Figure 2A). Using these reporters, neural crest cells not committed to the pigment cell lineage are *crestin:mCherry* single-positive, whereas *crestin:mCherry*/*mitfa:eGFP* double-positive cells are those newly committed to the pigment cell lineage. We treated embryos containing both reporter transgenes with BMPi from 12 to 24 HPF, during neural crest development and specification. At 24 HPF, we dissociated embryos and analyzed cells for fluorescent marker expression by flow cytometry (Figure 2A). Embryos treated with BMPi showed approximately a 1.5-fold increase in the percentage of *crestin:mCherry*/*mitfa:eGFP* double*-*positive cells per total *crestin:mCherry*-positive cells (Figure 2B, 2C). We further verified a change in specification by staining BMPi- or vehicle-treated *Tg(crestin:eGFP)* embryos with anti-Mitfa antibody and assessed the proportion of *crestin:eGFP*-positive cells that stained positive for Mitfa (Figure 2D). We observed a 1.3-fold increase in the proportion of Mitfa/*crestin:eGFP* double-positive cells per total *crestin:eGFP*-positive cells in animals treated with BMPi compared to vehicle control (Figure 2E). Altogether these results suggest that an increase in embryonic melanocytes is caused by an increase in the proportion of neural crest cells specified as pigment progenitor cells, rather than a change in proliferation of either neural crest or pigment progenitor cells.

**Figure 2.**
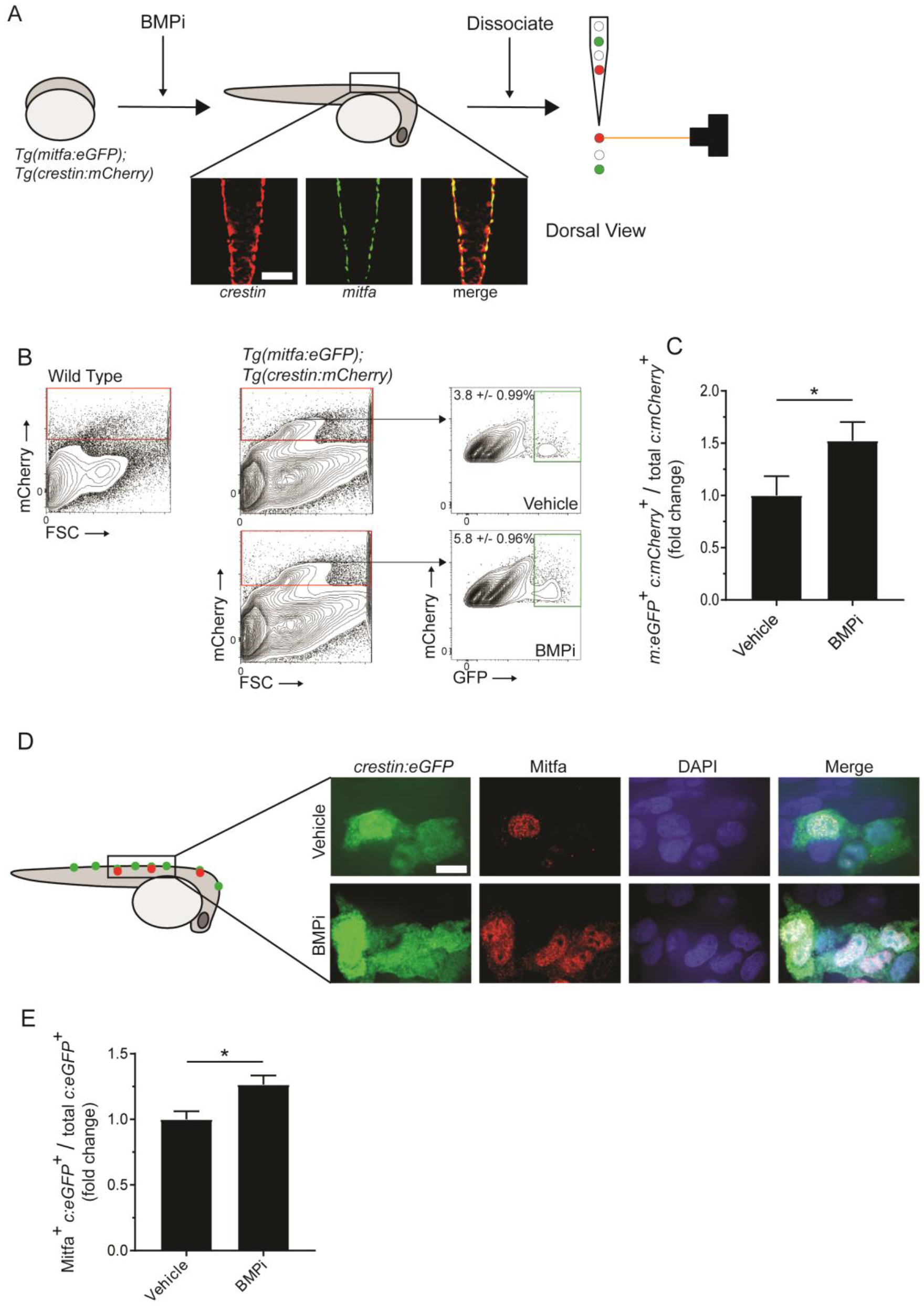
Inhibition of BMP signaling increases *mitfa*-positive neural crest cells. (A) Diagram of experiment. *Tg(crestin:mCherry); Tg(mitfa:eGFP)* embryos were treated with BMPi from 12 to 24 HPF. At 24 HPF, embryos were dissociated and analyzed via flow cytometry for GFP- and mCherry-positive cells, scale bar = 200 µm. (B) Gating strategy based on non-transgenic wild-type control to identify *crestin:mCherry*-positive cells and *crestin:mCherry/mitfa:eGFP* double-positive cells. Top, control vehicle-treated embryos. Bottom, BMPi-treated embryos. (C) Fold change in *crestin:mCherry/mitfa:eGFP* double-positive cells per total *crestin:mCherry-*positive cells in vehicle and BMPi-treated groups, n = 3 biological replicates of 80-100 stage-matched embryos pooled for each condition. *m:eGFP, mitfa:eGFP; c:mCherry, crestin:mCherry*. (D) anti-Mitfa immunofluorescence in *Tg(crestin:eGFP)* embryos treated with BMPi or vehicle control and fixed at 24 hours, scaled bar = 10 µm. (E) Fold change in Mitfa/*crestin:eGFP* double-positive cells per total *crestin:eGFP-*cells, n = 16 embryos for each condition. *c:eGFP, crestin:eGFP*. Error bars represent mean +/-SEM; P-value was calculated using ratio-paired t-test in panel C and Student’s t-test in panel E, * P < 0.05.

### BMP signaling in *mitfa-*expressing pigment progenitor cells can alter melanocyte development in embryogenesis

Because we observed an impact of BMP signaling on neural crest-to-pigment progenitor cell specification, we explored the relationship between *gdf6a* and *mitfa* expression. We performed *in situ* hybridization for *gdf6a* and *mitfa* during the course of neural crest and melanocyte development (Figure 3A). As described previously (Reichert et al., 2013; Rissi, Wittbrodt, Délot, Naegeli, & Rosa, 1995), *gdf6a* is expressed in the neural crest during induction. We observe downregulation of *gdf6a* in a rostrocaudal fashion as development proceeds. *mitfa* is expressed inversely, being turned on in neural crest cells rostrocaudally in the zone vacated by *gdf6a*. The timing of *gdf6a* and *mitfa* expression changes is consistent with the possibility that *gdf6a*-driven BMP signaling acts in neural crest cells to repress *mitfa* expression and prevent excess pigment progenitor cells from being specified. However, we also considered the possibility that BMP signaling is active in *mitfa*-positive cells and affects the fates of these cells. To determine if BMP signaling is active in *mitfa*-positive cells, we stained *Tg(mitfa:eGFP)* zebrafish with antibodies against phosphorylated-SMAD-1/5/8 (pSMAD). We verified specificity of the anti-pSMAD antibody using BMPi treated embryos (Figure S3). 30% of *mitfa*-expressing cells on the leading, caudal edge of the *mitfa* expression domain had nuclear-localized pSMAD staining, whereas only 7% of *mitfa-*expressing cells in regions rostral to the leading edge showed nuclear pSMAD staining (Figure 3B and 3C). These results suggest BMP signaling is active as *mitfa-*expressing cells first arise in the neural crest, but is turned off in such cells as development proceeds. To assess if BMP activity in *mitfa-*expressing cells can impact melanocyte development, we directly altered BMP activity in these cells. We first generated a stably transgenic zebrafish line expressing *gdf6a* under the control of the *mitfa* promoter *(Tg(mitfa:gdf6a))* to increase *gdf6a* expression in *mitfa-*expressing cells. Embryos expressing the *Tg(mitfa:gdf6a)* transgene developed fewer melanocytes than non-transgenic sibling controls (Figure 4A). To alter BMP signaling in a cell-autonomous manner within *mitfa-*expressing cells, we used the miniCoopR system in two complementary approaches: a) to express a dominant negative BMP receptor (dnBMPR), which suppresses intracellular BMP activity, and b) to express a phospho-mimetic variant of SMAD1 (SMAD1-DVD) to constitutively activate intracellular BMP activity (Ceol et al., 2011; Nojima et al., 2010; Pyati, Webb, & Kimelman, 2005). We injected *mitfa(lf)* animals with miniCoopR-dnBMPR, miniCoopR-SMAD1-DVD, or control miniCoopR-eGFP (Figure 4B). At 5 DPF, we scored animals for rescue of melanocytes. Animals injected with miniCoopR-dnBMPR showed a rescue rate of 79% as compared to 29% of miniCoopR-eGFP-injected animals. Furthermore, animals injected with miniCoopR-SMAD1-DVD showed a 15% rescue rate (Figure 4C). Together these results suggest BMP signaling is active in *mitfa-*expressing cells and modulating BMP signaling can alter the fate of these *mitfa*-expressing cells during development. Thus, *gdf6a*-driven BMP signaling can both limit the number of *mitfa*-expressing cells arising from the neural crest but also act in *mitfa*-expressing pigment progenitor cells to influence their development into melanocytes.

**Figure 3.**
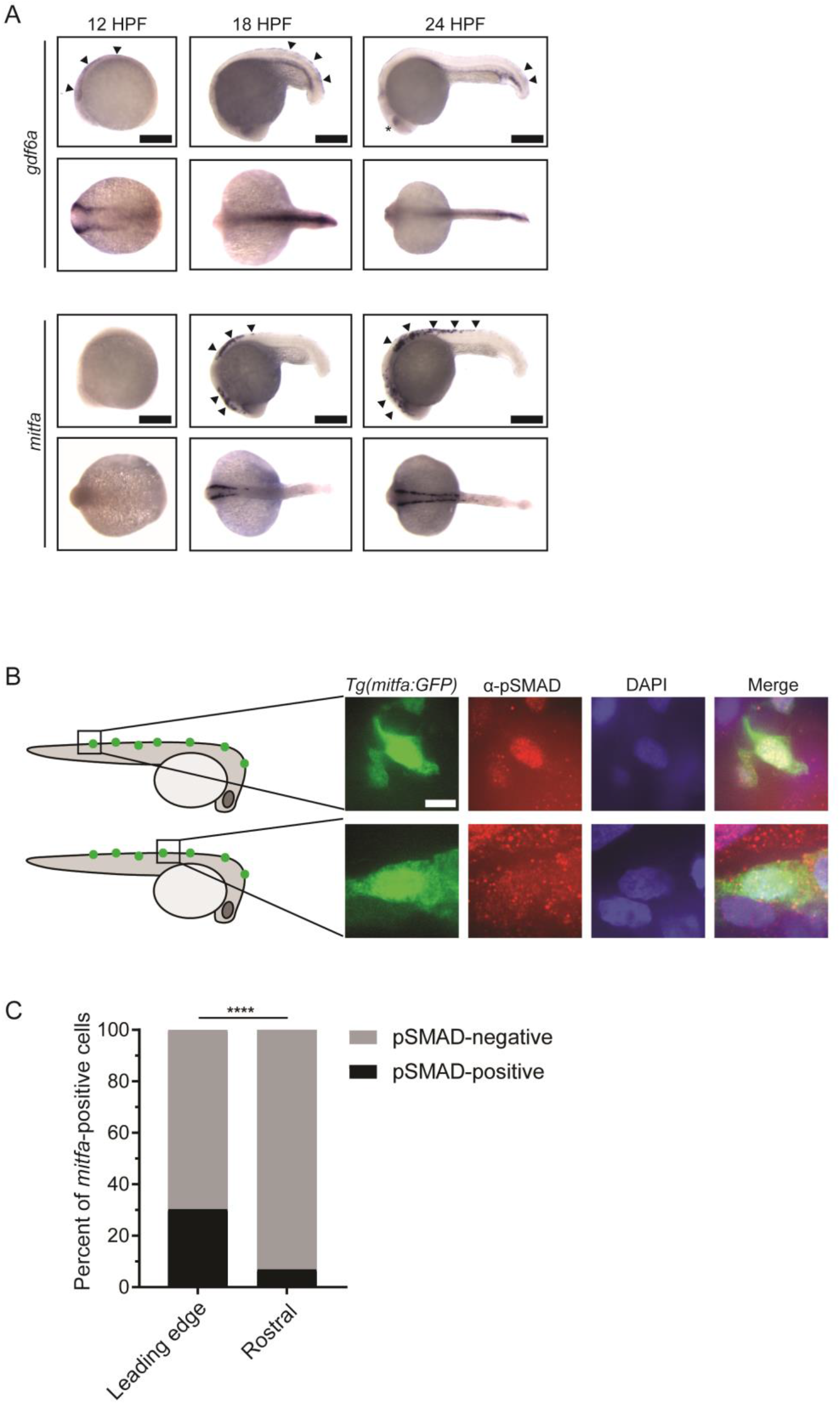
*gdf6a* expression and BMP activity in pigment progenitor cells. (A) RNA *in situ* hybridization for *gdf6a* (top) and *mitfa* (bottom) at 12-, 18-, and 24-hours post-fertilization. Arrowheads indicate expression domains in the neural crest of *gdf6a* and *mitfa*. Asterisk indicates known dorsal retinal expression of *gdf6a*. Scale bar = 500 µm. (B) Images of GFP-positive cells from *Tg(mitfa:eGFP)* zebrafish stained with α-pSMAD 1/5/8 antibody. Scale bar = 10 µm. (C) Quantification of *mitfa:eGFP-*positive cells that are phospho-SMAD1/5/8-positive. The leading edge encompassed 5 most caudal *mitfa*-positive cells, whereas rostral cells included any *mitfa*-positive cells rostral to the leading edge. n = 102 and 186 for distal leading edge and rostral cells, respectively. P-value was calculated using Fisher’s exact test, **** P < 0.0001.

**Figure 4.**
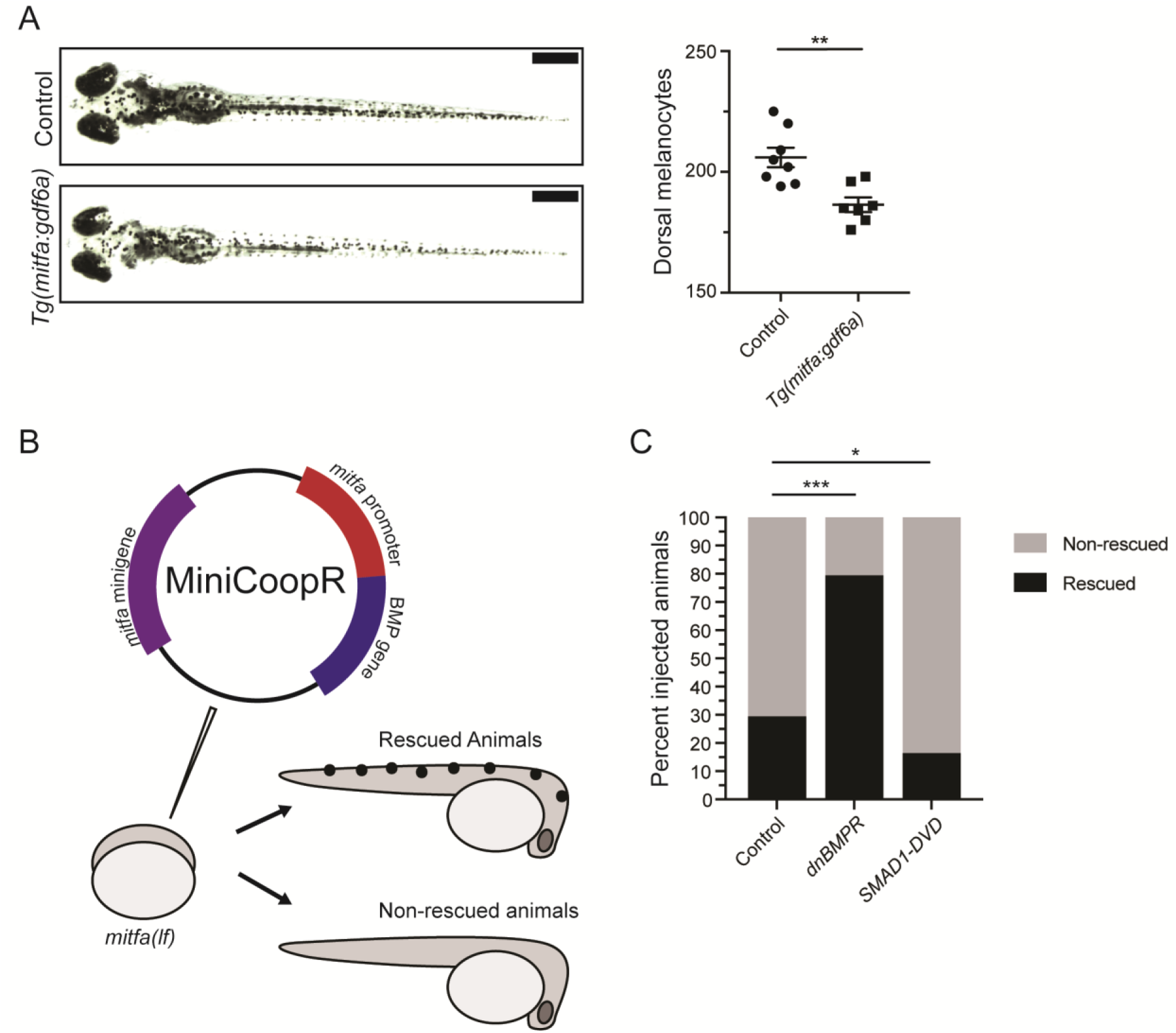
BMP signaling within pigment progenitor cells can impact embryonic melanocytes. (A) *Tg(mitfa:gdf6a)* and non-transgenic sibling control embryos (left), and quantification of dorsal melanocytes per animal in each group (right). Animals were treated with epinephrine prior to imaging at 5 DPF, n = 8 and 7 for control and *Tg(mitfa:gdf6a)* groups, respectively. Scale bar = 1 mm. (B) Diagram of miniCoopR rescue experiment. Animals harboring a *mitfa(lf)* mutation were injected at the single-cell stage with the miniCoopR vector containing a BMP gene. Animals were evaluated at 5 DPF for the presence of melanocytes. If melanocytes were present, that animal was scored as rescued, whereas animals lacking melanocytes were scored as non-rescued. (C) Percentages of rescued and non-rescued animals following injection of a miniCoopR-BMP vector, n = 361, 193 and 152 for control, *dnBMPR*, and *SMAD1-DVD* groups, respectively. Error bars represent mean +/-SEM. P-values were calculated Student’s t-test for panel A and with Fisher’s exact test with Bonferroni’s correction for panel C, * P <0.05, ** P <0.01, *** P <0.001.

### Iridophores, but not other neural crest derivatives, are reduced upon *gdf6a* loss

Because we observed no change in proliferation of *crestin*- or *mitfa*-positive populations, but the number of melanocytes developing from these precursors was increased, we questioned whether this increase corresponded with a commensurate loss of a related pigment or other neural crest-derived cell type. To determine what cells may be impacted, we looked for transcriptional changes in markers of other, related neural crest derivatives. We isolated RNA from *gdf6a(lf)* and wild-type embryos at 5 DPF. Additionally, we isolated RNA from embryos treated with a BMPi or vehicle control. We performed qPCR for markers of neural crest derivatives, including *mbpa* for glial cells, *pomca* for adrenal medullary cells, *neurog1* for neuronal cells, *aox5* for xanthophores, and *pnp4a* for iridophores (Fadeev et al., 2016; McGraw, Nechiporuk, & Raible, 2008; D.M. Parichy, Ransom, Paw, Zon, & Johnson, 2000; Thomas & Erickson, 2009). As a control, we used a chondrocyte marker, *col2a1a*, as craniofacial development has previously been described to be disrupted by *gdf6a* loss (Reed & Mortlock, 2010). Per our previous analysis, *gdf6a(lf)* mutants demonstrated an increase in expression of the melanocyte marker *tyrp1b* (Figure 1E). And as predicted based on previous literature, *gdf6a(lf)* mutants showed a decrease in expression of the chondrocyte marker, *col2a1a*. Markers for neuronal, glial, adrenal medullary, and xanthophore lineages were no different in *gdf6a* compared to wild-type animals (Figure 5A). Similar results were obtained in animals treated with a BMPi, with the exception of a change in *mbpa* expression, a marker for glial cells. Previous studies have shown glial cell development is regulated in part by BMP activity (Jin et al., 2001). Since *mbpa* expression was unchanged in *gdf6a(lf)* animals, this suggests another BMP ligand is involved in activating BMP signaling to promote glial cell development. For neuronal and xanthophore cell populations, we verified that the expression profile correlated with cell numbers or development of key structures. We treated animals with BMPi or vehicle and stained with anti-HuC/D antibody to label neuronal cells in the dorsal root ganglia and developing gastrointestinal tract (Lister et al., 2006) (Figure S4A-C). We detected no difference in dorsal root ganglia and enteric neuron development between each group. We imaged animals stably expressing *Tg(aox5:PALM-eGFP)* to label xanthophores and found no qualitative difference in xanthophores between BMPi- and vehicle-treated groups (Eom & Parichy, 2017) (Figure S4D). In our transcriptional analyses of *gdf6a(lf)* and BMPi-treated embryos, we observed a decrease in expression of *pnp4a*, a marker for the iridophore lineage, indicating a potential deficit of iridophore development (Figure 5A). Since *pnp4a* is expressed in other developing cells and tissues, such as retinal cell populations, we wanted to confirm these changes were specific to a deficit in neural crest-derived body iridophores (Cechmanek & McFarlane, 2017; Lopes et al., 2008; Petratou et al., 2018). We quantified the number of dorsal iridophores that developed in *gdf6a(lf)* embryos (Figure 5B) and embryos treated with BMPi (Figure 5C) at 5 DPF, using incident light to highlight embryonic iridophores. Embryos developed 32% and 27% fewer iridophores with *gdf6a(lf)* or BMPi treatment, respectively. Together, these results indicate that *gdf6a*-driven BMP signaling promotes iridophore development.

**Figure 5.**
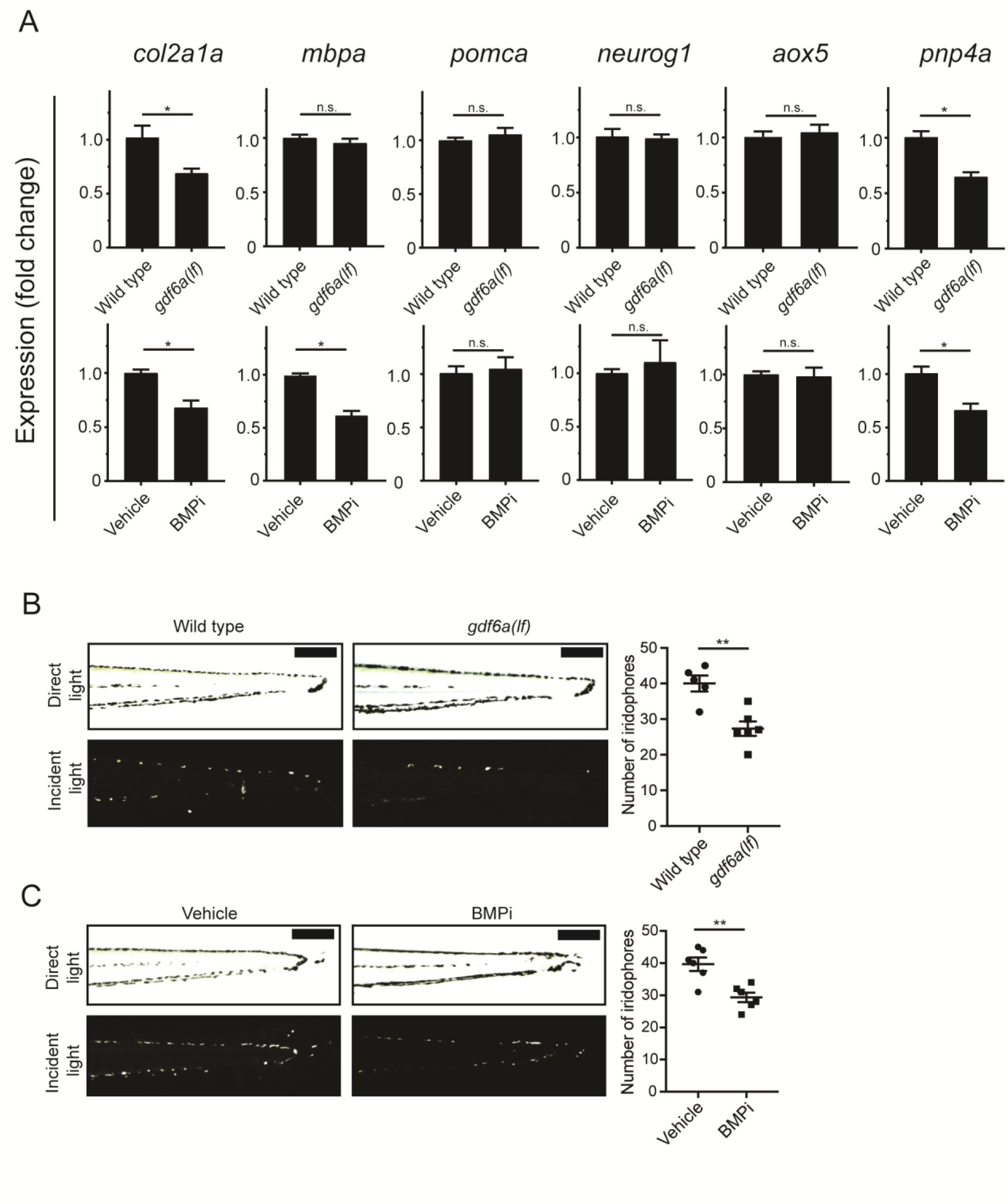
*gdf6a* loss and BMP inhibition impact development of specific neural crest derivatives. (A) Expression analyses of multiple neural crest lineage markers. qPCR was used to assess changes in markers of neural crest derivatives in *gdf6a(lf)* embryonic zebrafish (top) and BMPi-treated wild-type zebrafish (bottom) at 5 DPF; *col2a1a,* chondrocytes; *mbpa*, glial; *pomca*, adrenal medullary cells; *neurog1*, neuronal cells; *aox5*, xanthophores; *pnp4a*, iridophores; n = 5-6 for each group. (B) Direct light (top) and incident light (bottom) images of wild-type and *gdf6a(lf)* embryos at 5 DPF and quantification of dorsal iridophores (right) per animal in each group. Animals were treated with epinephrine prior to imaging at 5 DPF; n = 5 and 6 for wild-type and *gdf6a(lf)* groups, respectively; scale bar = 500 µm. (C) Direct light, top, and incident light, bottom, images of wild-type embryos treated with vehicle or BMPi from 12 to 24 HPF and quantification of dorsal iridophores, right, per animal in vehicle and BMPi treated groups. Animals treated with epinephrine prior to imaging at 5 DPF, n = 6 and 6 for vehicle and BMPi groups, respectively; scale bar = 1 mm. Error bars represent mean +/-SEM, P-values calculated with Student’s t-test, * P < 0.05, ** P < 0.01, n.s., not significant.

### BMP inhibition increases the likelihood a multipotent precursor will develop into a melanocyte

Melanocytes and iridophores have previously been shown to develop from *mitfa-*expressing pigment progenitor cells (Curran et al., 2010; Curran et al., 2009). To determine if BMP signaling regulates fate specification of melanocytes and iridophores from *mitfa*-expressing pigment progenitor cells, we performed lineage tracing. We injected *Tg(ubi:switch)* embryos, which stably express a *ubi:loxp-GFP-STOP-loxp-mCherry-STOP* transgene (Mosimann et al., 2011) with a *mitfa:Cre-ERT2* transgene to generate mosaic expression of Cre-ERT2 in *mitfa*-positive cells (Figure 6A). Injected embryos were treated with BMPi and hydroxytamoxifen (4-OHT), the latter to allow nuclear localization of Cre and generate recombinant events in individual *mitfa*-expressing pigment progenitor cells. Since these *mitfa*-expressing pigment progenitor cells are transient, 4-OHT treatment was limited to 12 to 24 HPF, with thorough fish water exchange to wash out the drug and prevent recombinant events after specification. At 5 DPF, embryos with individual recombinant events, indicated by single mCherry-positive cells, were evaluated for the fate of those cells. In animals treated with BMPi, we observed an increase in the ratio of labeled melanocytes to iridophores as compared to vehicle-treated controls (Figure 6B, S5). This result suggests that BMP signaling normally promotes the development of *mitfa*-expressing pigment progenitor cells into iridophores at the expense of melanocytes.

**Figure 6.**
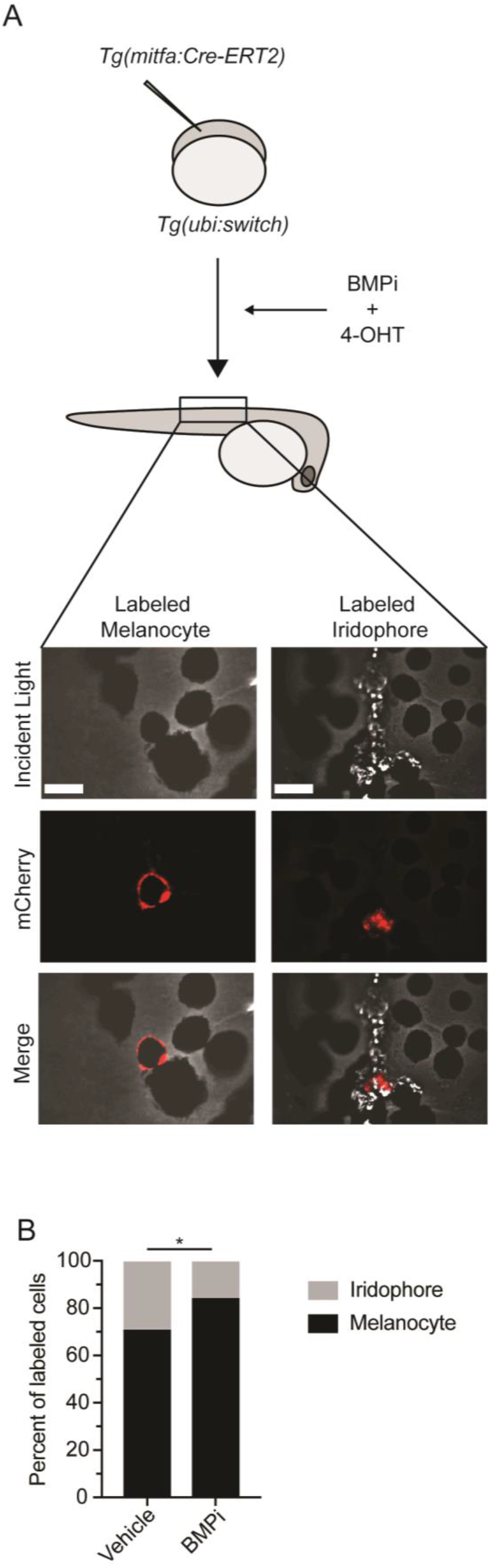
BMP inhibition impacts fate specification of *mitfa-*positive pigment progenitor cells. (A) Diagram of lineage tracing experiment. Embryos containing *Tg(ubi:switch)* were injected with a *mitfa:Cre-ERT2* construct and treated with BMPi and tamoxifen (4-OHT) from 12 to 24 HPF to block BMP signaling and allow Cre recombination. At 5 DPF, animals were screened for successful recombination by presence of single mCherry-labeled pigment cells, and the identities of those cells were assessed using incident light. Scale bar = 40 µm. (B) Quantification of mCherry-labeled cell fates at 5 DPF in vehicle and BMPi-treated animals, n = 101 and 80 for vehicle and BMPi groups, respectively; P-value calculated using Fisher’s exact test, * P < 0.05.

### BMP signaling represses *mitfa* expression within neural crest and pigment progenitor cells

Previous studies have indicated that the expression level of *mitfa* within pigment progenitor cells is important in specifying a melanocyte versus iridophore fate (Curran et al., 2010; Curran et al., 2009). Cells with a higher level *mitfa* expression are more likely to become melanocytes, while those that downregulate *mitfa* are more likely to become iridophores. Since *gdf6a(lf)* and BMP-inhibited embryos have excess melanocytes and fewer iridophores, we hypothesized that this phenotype resulted from disrupted regulation of *mitfa* expression in these embryos. This hypothesis was driven, in part, by our previous data in human melanoma cells, in which knockdown of *GDF6* decreased phospho-SMAD1/5/8 binding at the *MITF* promoter and increased *MITF* expression (Venkatesan et al., 2018). To assess *mitfa* levels within neural crest cells and *mitfa*-expressing pigment progenitor cells, we treated *Tg(crestin:eGFP)* and *Tg(mitfa:eGFP)* embryos with BMPi as previously described. We dissociated embryos and used fluorescence-activated cell sorting (FACS) to isolate *crestin-eGFP*-positive or *mitfa:eGFP-*positive cells. We then assessed *mitfa* transcript levels in each population by qPCR. Treatment with BMPi led to approximately 3-fold and 6-fold increases in *mitfa* expression in *crestin:eGFP*-positive and *mitfa:eGFP-*positive cells, respectively (Figure 7A). To explore this question on a single-cell level and analyze Mitfa protein levels, we stained BMPi-treated and vehicle-treated *Tg(crestin:eGFP)* embryos with an anti-Mitfa antibody (Figure 7B) (Venkatesan et al., 2018). In BMPi-treated animals, we observed a 2.5-increase in Mitfa staining intensity in *crestin:eGFP-*positive cells, indicating inhibition of BMP signaling leads to an increase in Mitfa protein in pigment progenitor cells at a single-cell level (Figure 7C). Furthermore, those cells that were Mitfa-positive and *crestin:eGFP-*negative showed a 1.7-fold increase in Mitfa staining intensity, indicating inhibition of BMP signaling also leads to an increase in Mitfa protein following specification of pigment cells (Figure 7B, 7C). Together, these results indicate BMP signaling suppresses *mitfa* expression in cells during specification of pigment cell lineages.

**Figure 7.**
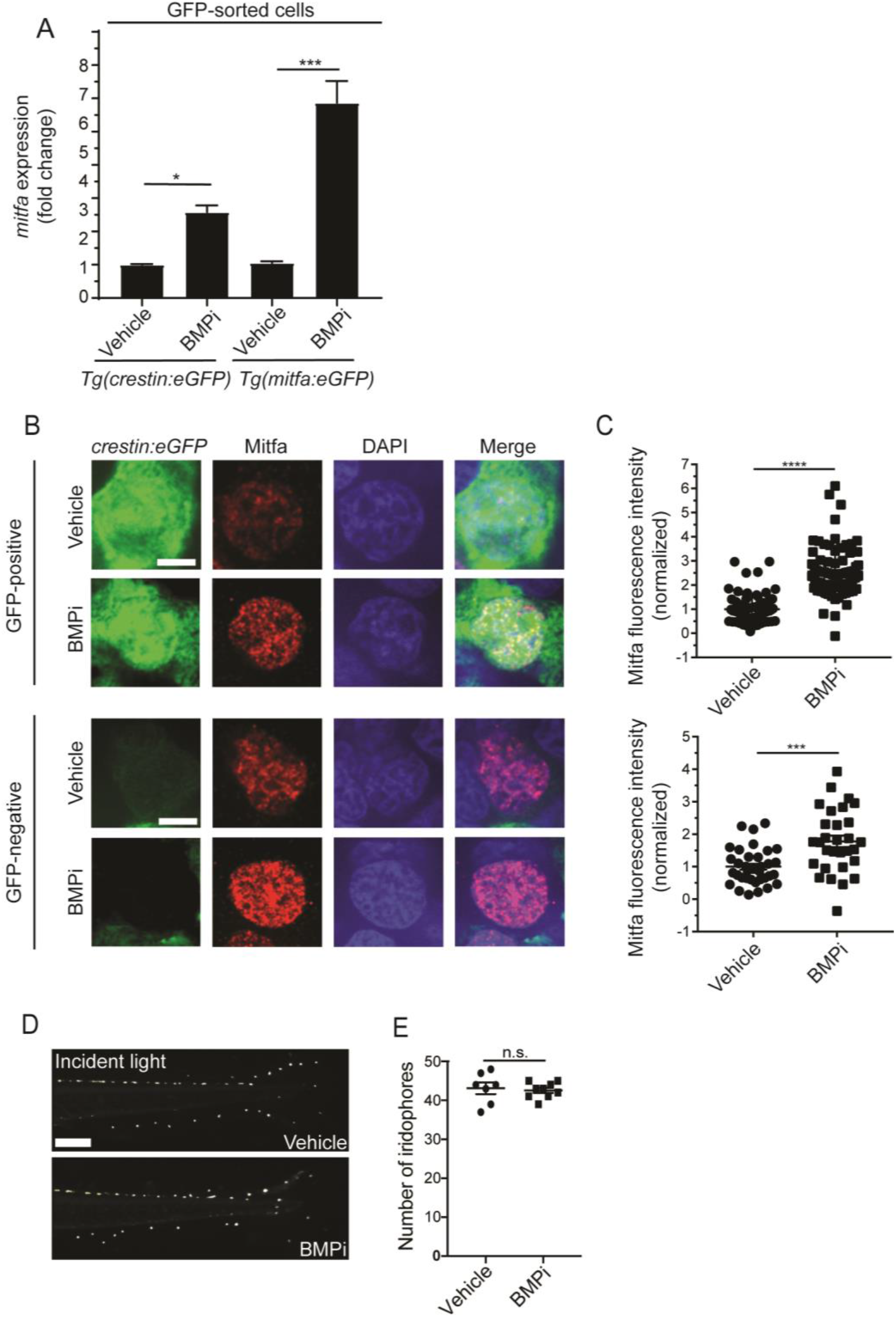
BMP signaling regulates expression of and acts through *mitfa* to impact pigment cell fates. (A) *mitfa* expression in sorted GFP-positive cells from *Tg(crestin:eGFP)* and *Tg(mitfa:eGFP)* embryos treated with vehicle or BMPi from 12 to 24 HPF, n = 4-5 for each condition. (B) anti-Mitfa immunofluorescence, DAPI and merged images of *Tg(crestin:eGFP)* embryos treated with vehicle control or BMPi in GFP-positive cells (top) and GFP-negative cells (bottom), scale bar = 5 µm. (C) Quantification of anti-Mitfa fluorescence intensity of individual nuclei in GFP-positive cells (top) and GFP-negative cells (bottom); n = 65 and 74 for GFP-positive vehicle and BMPi groups, respectively; n = 35 and 30 for GFP-negative vehicle and BMPi groups, respectively. (D) Incident light images of *mitfa(lf)* embryonic zebrafish treated with vehicle or BMPi from 12 to 24 HPF and imaged at 5 DPF, scale bar = 1 mm. (E) Quantification of dorsal iridophores in *mitfa(lf)* embryonic zebrafish treated with vehicle or BMPi from 12 to 24 HPF, n = 7 and 9 for vehicle and BMPi groups, respectively. Error bars represent mean +/-SEM, P-value was calculated using one-way ANOVA with Tukey’s multiple comparisons test in panel A and Student’s t-test in panel C and E. * P <0.05, *** P <0.001, **** P<0.0001, n.s., not significant.

### Regulation of pigment cell fate by BMP signaling is dependent on *mitfa*

If deregulated *mitfa* expression is critical to the phenotypic defects observed upon inhibition of BMP signaling, then these defects should be dependent on *mitfa* function. To determine whether *mitfa* is indeed responsible for mediating the shift in cell fate regulated by BMP activity, we treated *mitfa(lf)* embryos with BMPi. As *mitfa* is necessary for the specification of all body melanocytes, *mitfa(lf)* animals do not develop any melanocytes during embryogenesis or through adulthood. However, these animals can develop iridophores and develop a greater number of iridophores at baseline than their wild-type counterparts (Lister et al., 1999). We hypothesized that, if an elevation of *mitfa* expression in BMPi-treated embryos was required to shift pigment progenitor cell fates from iridophores to melanocytes, there would be no decrease in the number of iridophores when *mitfa(lf*) embryos were treated with BMPi. Indeed, BMPi-treated embryos showed no difference in the number of iridophores compared to vehicle-treated controls (Figure 7D, 7E). Together, these results indicate that BMP inhibition requires *mitfa* to direct pigment progenitor cells away from iridophore fate.

## Discussion

Our results elucidate a role for *gdf6a*-activated BMP signaling in suppressing melanocyte development from the neural crest during embryogenesis. Inhibition of BMP signaling leads to an increase of neural crest cells expressing *mitfa*, affecting the proportion of neural crest cells specified as pigment progenitor cells. Additionally, in BMP-inhibited embryos these *mitfa-*positive pigment progenitor cells demonstrate an increased propensity to become melanocytes, instead of iridophores. Cells in BMP-inhibited embryos have increased expression of *mitfa*, and the function of *mitfa* is required for the reduction of iridophores observed in BMP-inhibited embryos. Based on these findings, we propose that *gdf6a*-activated BMP signaling normally represses *mitfa* expression, limiting both the development of pigment progenitor cells from the neural crest and the specification of melanocytes from these pigment progenitor cells. As discussed below, *MITF* is downregulated by *GDF6*-activated BMP signaling to prevent melanocytic differentiation in melanomas (Venkatesan et al., 2018). The function we have defined for *gdf6a*-activated BMP signaling in development suggests that its activity is co-opted in tumors to prevent differentiation of melanoma cells.

### Regulation of pigment cell fate by BMP signaling

Our studies indicate *gdf6a*-activated BMP signaling can regulate pigment cell development from the neural crest in two ways. First, BMP signaling restricts the number of neural crest cells that transition into *mitfa*-positive pigment cell progenitors. When BMP signaling is abrogated, additional cells adopt a pigment progenitor fate, which likely is a source of supernumerary melanocytes. Second, BMP signaling biases the fate choice of *mitfa*-positive progenitor cells. In BMP-deficient embryos, *mitfa*-positive progenitor cells more often become melanocytes and less often become iridophores. Previous studies have suggested a common melanocyte-iridophore progenitor (Curran et al., 2010; Curran et al., 2009; Petratou et al., 2018), and our data support the existence of such a progenitor and indicate that it is *mitfa*-expressing and influenced by BMP signaling. While BMP signaling regulates the fate of a common melanocyte-iridophore precursor, the decrease in the number of iridophores cannot fully account for the number of melanocytes gained in *gdf6a(lf)* and BMPi-treated embryos. Because *gdf6a(lf)* and BMPi-treatment are potentially impacting the entirety of neural crest development, other neural crest cells may be mis-specified to the melanocyte lineage. This misspecification could account for the discrepancy between the gain of melanocytes and loss of iridophores. If misspecification of other neural crest cells is occurring, other neural crest lineages could show a deficit. However, in our assays evaluating other lineages, we detected no deficits outside of a loss of iridophores. Among several possibilities, the deficit may be present in a neural crest lineage we did not directly measure. Alternatively, deficits in other neural crest lineages may be small and distributed across multiple other lineages, such that our assays are unable to detect those subtle changes. Lastly, proliferation within the neural crest and of neural crest derivatives following migration from the crest is known to occur (Dougherty et al., 2013; Gianino, Grider, Cresswell, Enomoto, & Heuckeroth, 2003), and it is possible that such proliferation could compensate for any deficit. In summary, the supernumerary melanocytes observed in *gdf6a(lf)* and BMPi-treated embryos are likely to arise from some combination of neural crest cells that are shunted to the pigment cell lineage and melanocyte-iridophore precursors that preferentially adopt a melanocyte fate.

### Regulation of *mitfa* by BMP signaling

Our studies identify *gdf6a*-activated BMP signaling as a regulator of *mitfa* during pigment cell development in zebrafish. Previous studies have identified roles for *gdf6a* in the preplacodal ectoderm, retinal cell survival, and craniofacial development in zebrafish, while others have broadly connected BMP signaling to fate determination and cell survival in the neural crest in other model systems (French, Erickson, French, Pilgrim, & Waskiewicz, 2009; Gosse & Baier, 2009; Hanel & Hensey, 2006; Jin et al., 2001; Reed & Mortlock, 2010; Reichert et al., 2013). However, the specific role of BMP signaling and of *gdf6a* on pigment cell development has heretofore been uncharacterized. Our analyses indicate that *gdf6a* is expressed in neural crest cells prior to the rostrocaudal onset of *mitfa* expression. In addition, we observed an overlap of BMP activity and *mitfa* expression at the leading edge of the rostrocaudal *mitfa* progression. When BMP signaling was inhibited, we found increased expression in neural crest cells of *mitfa* RNA and Mitfa protein. Together, these results suggest that *gdf6a*-driven BMP signaling regulates expression of *mitfa* and, consequently, directs fates adopted by *mitfa*-expressing cells. We speculate that such a role underlies the excess melanocytes observed in *gdf6a(lf)* and BMPi-treated embryos. In the absence of *gdf6a* and BMP signaling, increased expression of *mitfa* could lead to a greater proportion of neural crest cells adopting a pigment cell fate and could lead to a greater propensity of melanocyte-iridophore precursors adopting a melanocyte fate. These findings are consistent with what has previously been established in human melanoma cells, where *GDF6-*activated BMP signaling has been shown to promote pSMAD binding to *MITF* and is suspected to directly regulate *MITF* expression (Venkatesan et al., 2018). Our results support this regulatory role and provide a developmental context *in vivo* to understand why *GDF6*-activated BMP signaling is able to regulate *MITF* in melanoma cells.

### Reiteration of normal physiologic function in melanoma

*GDF6* and BMP signaling were previously described in melanoma to suppress differentiation through binding of pSMAD to *MITF* and corresponding repression of *MITF* expression (Venkatesan et al., 2018). Results from the current study indicate *gdf6a* and BMP signaling likely act in a similar fashion during development to repress expression of *MITF*, either directly or indirectly, leading to suppression of melanocyte specification and differentiation from the neural crest. Together, these findings suggest BMP activity in melanoma is a recapitulation of normal regulatory functions executed by *gdf6a* and BMP signaling during pigment cell development. It has been previously established that lineage programs can be co-opted by cancers to promote pro-tumorigenic characteristics (Carreira et al., 2006; Gupta et al., 2005). These programs activate EMT factors, such as *TWIST1* and *SNAI2*, and factors associated with neural crest multipotency, such as *SOX10*, to promote invasiveness, proliferative capacity, metastatic capability, and therapeutic resistance (Caramel et al., 2013; Casas et al., 2011; Shakhova et al., 2015). However, it is unclear if these factors have similar regulation between normal development and melanoma. Here, we have described a developmental role for *GDF6* that is reiterated in a pathologic process in disease. Because initiation and maintenance of neural crest gene expression has been shown to be important in melanoma, a better understanding of how regulation occurs during development may have clinical implications (Kaufman et al., 2016). Our findings indicate BMP signaling has a regulatory role over key differentiation genes during melanocyte development from the neural crest. Many studies have implicated expression of neural crest and melanocyte factors during many phases of melanoma, including initiation, progression, invasion, metastasis, and therapeutic resistance of melanoma (Carreira et al., 2006; Fallahi-Sichani et al., 2017; Gupta et al., 2005; Kaufman et al., 2016; Shaffer et al., 2017). Taken together, these findings suggest therapeutic targeting of GDF6 or BMP signaling would likely have a positive impact on prognosis and outcome in melanoma patients by promoting differentiation in tumors.

## Acknowledgements

We thank Nathan Lawson for pcsDest2 plasmid; David Kimelman for the dnBMPR plasmid; Takenobu Katagiri for the SMAD1-DVD plasmid; Christian Mossiman for *Tg(ubi:switch)* zebrafish strain and CreERT2 plasmid; Charles Kaufmann for *Tg(crestin:eGFP)* and *Tg(crestin:mCherry)* zebrafish strains; Patrick White, Ed Jaskolski and the staff at the UMMS Animal Medicine Department for fish care; Tammy Krumpoch for guidance and assistance in performing flow cytometry and FACS experiments. AKG was supported by a Melanoma Research Foundation Looney Legacy Foundation Medical Student Award, Center for Translational Sciences TL1 Training Fellowship through the UMMS CCTS (UL1-TR001453), NCI NRSA F31 Predoctoral Fellowship (1F31CA239478-01). Research was supported by a Kimmel Scholar Award (SKF-13-123), Department of Defense Peer Reviewed Cancer Research Program Career Development Award (W8IXWH-13-0107) and NIH National Institute of Arthritis and Musculoskeletal and Skin Diseases grant (R01AR063850) to CJC. The content is solely the responsibility of the authors and does not necessarily represent the official views of the Department of Defense or NIH.

## Author Contributions

AKG, AMV, and CJC conceived the project. AKG, AMV, and CJC designed and interpreted the melanocyte quantification experiments and results. AKG and CJC designed and interpreted the proliferation, specification, and neural crest lineage experiments and results. AKG performed the flow cytometry, qPCR, immunofluorescence, lineage tracing and *in situ* hybridization experiments. AKG and AMV performed the melanocyte quantification experiments. AMV generated the MiniCoopR and pCS-DEST plasmids and generated the probes for *in situ* hybridization. AMV and MG generated and isolated *gdf6b* mutant zebrafish. AKG and CJC wrote the manuscript.

## Methods

### Zebrafish

Zebrafish were handled in accordance with protocols approved by the University of Massachusetts Medical School IACUC. Fish stocks were maintained in an animal facility at 28.5°C on a 14-hour/10-hour Light/Dark cycle (Westerfield, 1995). The wild-type strain used was AB. Published strains used in this study include *gdf6a(lf) (gdf6a^s327^) (Gosse & Baier, 2009)*, *Tg(mitfa:eGFP)* (Curran et al., 2009), *Tg(crestin:eGFP)* (Kaufman et al., 2016), *Tg(crestin:mCherry) (Kaufman et al., 2016)*, *mitfa(lf)* (Lister et al., 1999), *Tg(ubi:switch)* (Mosimann et al., 2011), *Tg(aox5:PALM-eGFP)* (Eom & Parichy, 2017). Construction of new strains generated are detailed below.

### DNA Constructs

DNA constructs were built using Gateway cloning (Life Technologies). Sequences of *gdf6a*, *dnBMPR* (Pyati et al., 2005) and *SMAD1-DVD* (Nojima et al., 2010) were PCR-amplified and cloned into pDONR221 (Life Technologies). Oligonucleotides used in cloning are described in Key Resources Section. Previously published entry clones used in this study were pENTRP4P1r-*mitfa* (Ceol et al., 2011), pDONR221-*gdf6b* (Venkatesan et al., 2018), pDONR221-*CreERT2* (Mosimann et al., 2011). Previously published destination vectors used in this study are MiniCoopR (MCR) (Ceol et al., 2011) and pcsDest2 (Villefranc, Amigo, & Lawson, 2007). p3E-*polyA*, pME-*eGFP*, pDestTol2CG2, pDestTol2pA2, pCS2FA-transpoase were acquired from the Tol2Kit (Kwan et al., 2007). Using the entry clones and destination vectors described above, the following constructions were built using multisite or single site Gateway (Life Technologies): MCR-mitfa:dnBMPR:pA, MCR-mitfa:eGFP:pA, MCR-mitfa:SMAD1-DVD:pA, pDestTol2CG2-mitfa:gdf6a:pA, pDestTol2pA2-mitfa:CreERT2:pA, pcsDest2-gdf6a, pcsDest2-gdf6b. All constructs were verified by restriction digest or sequencing.

### Construction of *gdf6b(lf)*

To generate *gdf6b(lf)* mutants, we used TALEN genome editing. TALEN’s were designed targeting exon 1 of *gdf6b* (TAL1 sequence: GTCAGCATCACTGTTAT; TAL2 sequence: CCTTGATCGCCCTTCT). TALENs were assembled using the Golden Gate TALEN kit (Addgene) per the manufacturer’s instructions. TALEN plasmids were linearized and transcribed with mMESSAGE mMACHINE kit (Ambion). Zebrafish embryos were injected with 50 pg of mRNA of each TALEN arm. Injected embryos (F0) were matured to breeding age and outcrossed. Resulting offspring (F1) were genotyped by extraction genomic DNA from fin clips per standard protocol and PCR amplification with *gdf6b* primers. F1 offspring carrying mutations by genotyping were sequenced to identify mutations predicted to lead to loss of function of *gdf6b*. Following identification of candidate zebrafish by sequencing, zebrafish were bred to generate homozygous *gdf6b(lf)* mutations. Whole RNA was isolated from homozygous *gdf6b(lf)* embryos at 20 HPF and qPCR was used to determine effective depletion of *gdf6b* transcripts. Primers for genotyping and qPCR are listed in the Key Reagents section.

### Construction of *Tg(mitfa:gdf6a)*

To generate the *Tg(mitfa:gdf6a)* transgenic line, 25 pg of pDestTol2CG2-mitfa:gdf6a:pA was injected along with 25 pg of Tol2 transposase RNA, synthesized from pCS2FA-*transposase*, into single cell wild-type embryos (Kwan et al., 2007). Embryos were screened for incorporation of the transgene by expression of cmlc:eGFP in the heart at 48 HPF. Animals with eGFP-positive hearts (F0) were outcrossed to wild-type animals to determine germline incorporation.

### Drug Treatments

Drugs used in experiments were reconstituted at stock concentrations in solvent as follows: DMH1 (BMPi), 10mM in DMSO; Tamoxifen (4-OHT), 1mg/mL in ethanol; Epinephrine, 10mg/mL in embryo media. Embryos were dechorionated by incubating in Pronase (Roche) for 10 minutes with gentle shaking. Dechorionated embryos were transferred to 6-well plates coated in 1.5% agarose in embryo media. Embryo media with appropriate drug concentration or vehicle control was added to each well. For BMPi and 4-OHT treatments, embryos were treated from 12 HPF (6ss) to 24 HPF (Prim-5). Embryos were incubated at 28.5°C for the duration of the drug treatment. Following drug treatment, embryos were thoroughly washed in fresh embryo medium and returned to incubator in new embryo medium until analysis.

### Lineage Tracing

To trace the lineage of embryonic pigment cells, *Tg(ubi:switch)* embryos were injected with 25 pg of pDestTol2pA2-mitfa:Cre-ERT2:pA and 25 pg of Tol2 transposase RNA at the single-cell stage. At 12 HPF, injected embryos were treated with BMPi and 4-OHT as described above. Following treatment, embryos were thoroughly washed and allowed to mature at 28.5°C to 5 DPF. Embryos were treated with 1 mg/mL epinephrine to contract melanosomes, anesthetized using 0.17mg/mL tricaine in embryo media, mounted in 1% low-melt agarose on a plastic dish, and submerged in embryo media for imaging.

### Mosaic Rescue

MiniCoopR constructs MCR-mitfa:dnBMPR:pA, MCR-mitfa:SMAD1-DVD:pA, and MCR-mitfa:eGFP:pA (control) were used. *mitfa(lf)* animals were injected with 25 pg of a single construct and 25 pg of Tol2 transposase RNA. Upon successful integration of the MCR constructs, the *mitfa*-minigene in the construct allowed development of melanocytes. Embryos were screened for incorporation of the transgene by rescue of melanocytes at 5 DPF (Ceol et al., 2011).

### *In* Situ Hybridization

RNA sense and anti-sense probes were synthesized from pcsDest2-*gdf6a* and pcsDest2-*gdf6b* constructs using DIG RNA Labeling Kit (Roche) per the manufacturer’s instruction. Wild-type embryos of the appropriate stage were fixed in 4% PFA at 4°C for 24 hours. Following fixation, embryos were dehydrated in methanol at stored at −20°C. Whole mount *in situ* hybridization was performed as previously described (Reichert et al., 2013). Hybridized probes were detected using anti-digoxigenin (DIG) antibodies tagged with alkaline-phosphatase (AP) (Roche) using NBT/BCIP (Roche) solution per the manufacturer’s instructions. Stained embryos were mounted in 2.5% methylcellulose and imaged using a Leica M165FC microscope and Leica DFC400 camera. Specificity of the probes was verified using sense probes synthesized from the same construct.

### Immunofluorescence

Embryos were fixed at the desired time or following drug treatment in 4% PFA for 24 hours at 4°C. Whole mount immunofluorescence was performed as previously described (Venkatesan et al., 2018). Primary antibodies used were pSMAD-1/5/8 (1:100 dilution) (Cell Signal Technologies), HuC/D (1:100 dilution) (Sigma), mitfa (1:100 dilution) (Venkatesan et al., 2018). AlexaFluor-488 (Invitrogen) and AlexaFluor-555 (Invitrogen) conjugated secondary antibodies were used to detect primary antibody signaling. Nuclei were counterstained with DAPI. Following staining, animals were dissected to remove yolk sack and flat mounted laterally on slides using VectaShield mounting medium. Fluorescent images were taken using a Leica DM5500 microscope with a Leica DFC365FX camera, and a Zeiss Axiovert 200 microscope outfitted with a Yokogawa spinning disk confocal scanner. Cells and structures were counted, and data was analyzed using Microsoft Excel and GraphPad Prism 7.

### Flow Cytometry & Fluorescence Activated Cell Sorting (FACS)

Embryos were treated and matured to appropriate age as per drug treatment protocol described above. At a desired timepoint, embryos were washed in PBS and transferred to 500 µL of PBS + 5% FBS (FACS buffer). Embryos were mechanically dissociated in FACS buffer using a mortar and pestle. Dissociated embryos were washed with FACS buffer and filtered through a 40 µm mesh membrane. Samples were analyzed using a BD FACS Aria II flow cytometer and sorted directly into Trizol LS (Life Technologies) for RNA isolation. Flow cytometry data was analyzed using FlowJo software (Becton, Dickinson & Company) and GraphPad Prism 7.

### Quantitative Real-Time PCR (qPCR)

Oligos used for qPCR primers are listed in Key Reagents section. RNA was isolated from FACS-sorted cells or whole embryos using Trizol reagent (Life Technologies) and purified using the RNeasy kit (Quiagen) per manufacturer’s protocol. cDNA was synthesized from purified RNA using the SuperScript III First Strand Synthesis kit (Thermo Fisher). Reaction mixes were assembled with SYBR Green RT-PCR master mix (Thermo Fisher), primers, and 25 ng cDNA, and analyzed using a StepOnePlus Real Time PCR System (Applied Biosystems). Fold changes were calculated using the ΔΔCt method using Microsoft Excel and GraphPad Prism 7.

### Imaging and Quantification

Zebrafish adults and embryos were treated with 1 mg/mL epinephrine to contract melanosomes prior to imaging unless otherwise noted. Fish were anesthetized in 0.17% Tricaine in embryo media and positioned in 2.5% methylcellulose in embryo media for imaging. Images of adult fish were captured with a Nikon D90 DSLR camera. Brightfield and incident light images of embryos were captured with Leica M165FC microscope and Leica DFC400 camera. Fluorescent images of embryos were captured with a Leica DM5500 upright microscope with a Leica DFC365FX camera, and a Zeiss Axiovert 200 microscope outfitted with a Yokogawa spinning disk confocal scanner. Images were processed using ImageJ and Leica LAS X software. Cells were counted and analyses were performed using Microsoft Excel and GraphPad Prism 7. Statistical calculations were performed using GraphPad Prism 7 as described in each Figure legend.

### Statistical Analysis

Statistical analyses were performing using GraphPad Prism 7 software package. Statistical significance of experiments was calculated using Student’s t-test, ratio-paired t-test, Fisher’s exact test with Bonferroni’s correction, 1-way ANOVA with Tukey’s multiple comparison test as described in each figure legend. Statistical significance was denoted as follows: not significant (ns) P > 0.05, *P<0.05, **P<0.01, ***P<0.001 and ****P<0.0001.

---

**Figure S1 (Related to Figure 1).**
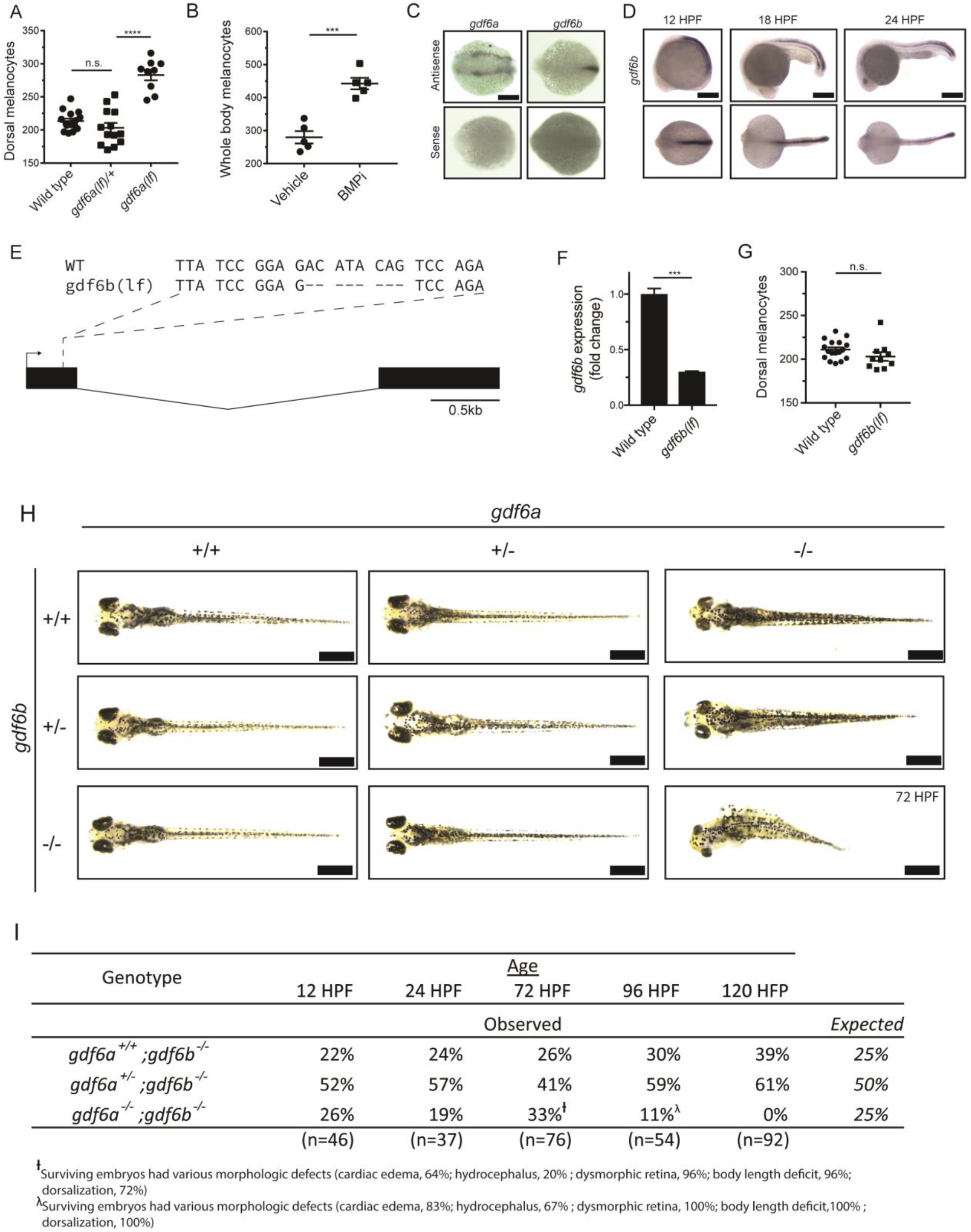
*gdf6* paralogs are necessary for normal embryonic development. (A) Quantification of dorsal melanocytes in *gdf6a(lf/+)* heterozygotes, *gdf6a(lf)* homozygotes and wild-type embryos. (B) Quantification of whole-body melanocytes in vehicle- and BMPi-treated embryos. (C) Verification of *gdf6a* and *gdf6b* probe specificity. (D) RNA *in situ* hybridization for *gdf6b* at 12-, 18-, and 24-hours post-fertilization, scale bar = 500 µm. (E) Sequence of *gdf6b(lf)* mutant indicating deletion and frameshift in exon 1. (F) Decreased *gdf6b* expression in *gdf6b(lf)* embryos. (G) Quantification of dorsal melanocytes in *gdf6b(lf)* mutants compared to wild-type embryos. (H) Images of *gdf6a(lf)* and *gdf6b(lf)* mutant combinations. *gdf6b(lf)* animals have no morphologic defects compared to wild-type embryos at 5 DPF, while *gdf6a(lf)* animals show pigmentation and eye morphology defects. *gdf6a(lf);gdf6b(lf)* double mutants show significant morphologic defects associated with *gdf6a(lf)* as well as decreased body length, cardiac edema and hydrocephalus. Scale bar = 1 mm. (I) Survival of *gdf6b(lf)* embryos with *gdf6a(lf)* mutations. ^†^, surviving embryos had various morphologic defects (cardiac edema, 64%; hydrocephalus, 20%; dysmorphic retina, 96%; body length deficit, 96%; dorsalization, 72%). λ, surviving embryos had various morphologic defects (cardiac edema, 83%; hydrocephalus, 67%; dysmorphic retina, 100%; body length deficit, 100%; dorsalization, 100%). Error bars represent mean +/-SEM. P-values were calculated using one-way ANOVA with Tukey’s multiple comparison test for panel A and with Student’s t-test for panels B, F, and G. *** P < 0.001, **** P <0.0001, n.s., not significant.

**Figure S2 (Related to Figure 2).**
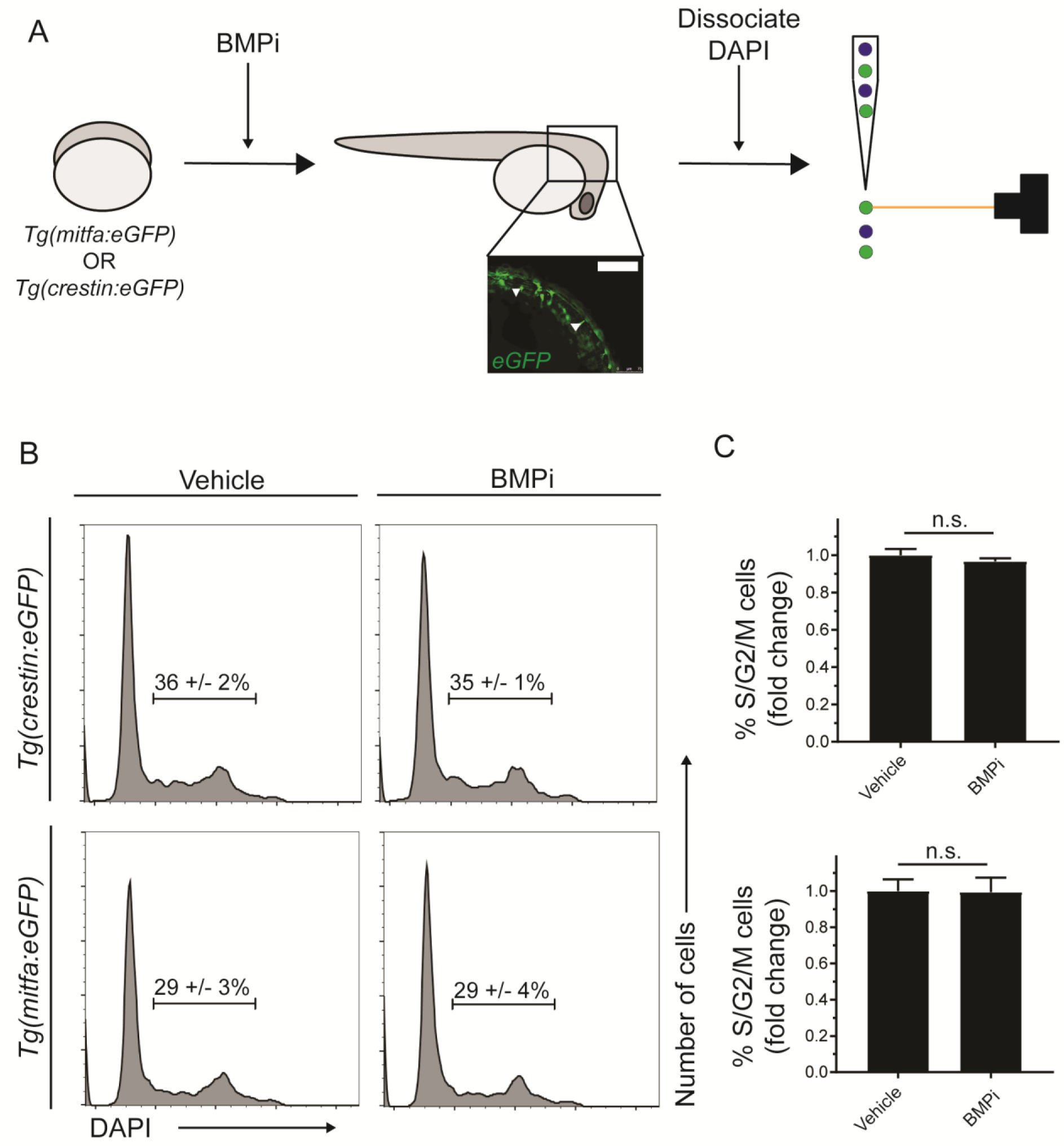
Increased proliferation is not observed in neural crest and pigment progenitor cell populations of BMPi-treated embryos. (A) Diagram of experiment. Embryos expressing either *Tg(mitfa:eGFP)* or *Tg(crestin:eGFP)* were treated with BMP inhibitor from 12 to 24 HPF. Following treatment, embryos were dissociated, fixed, and stained for DNA content using DAPI and analyzed via flow cytometry. Scale bar = 200 µm. (B) Flow cytometry histograms showing the percentage of cells in S/G2/M phases in *crestin:eGFP*-positive or *Tg(mitfa:eGFP*-positive cell populations in BMPi-treated embryos compared to vehicle-treated embryos. (C) Fold change of *crestin:eGFP*-positive cells (top) and *mitfa:eGFP*-positive cells (bottom) in S/G2/M phases. n = 4 biological replicates of 80-100 stage-matched embryos pooled for each condition. Error bars represent mean +/-SEM, P-value calculated using ratio paired t-test, n.s., not significant.

**Figure S3 (Related to Figure 3).**
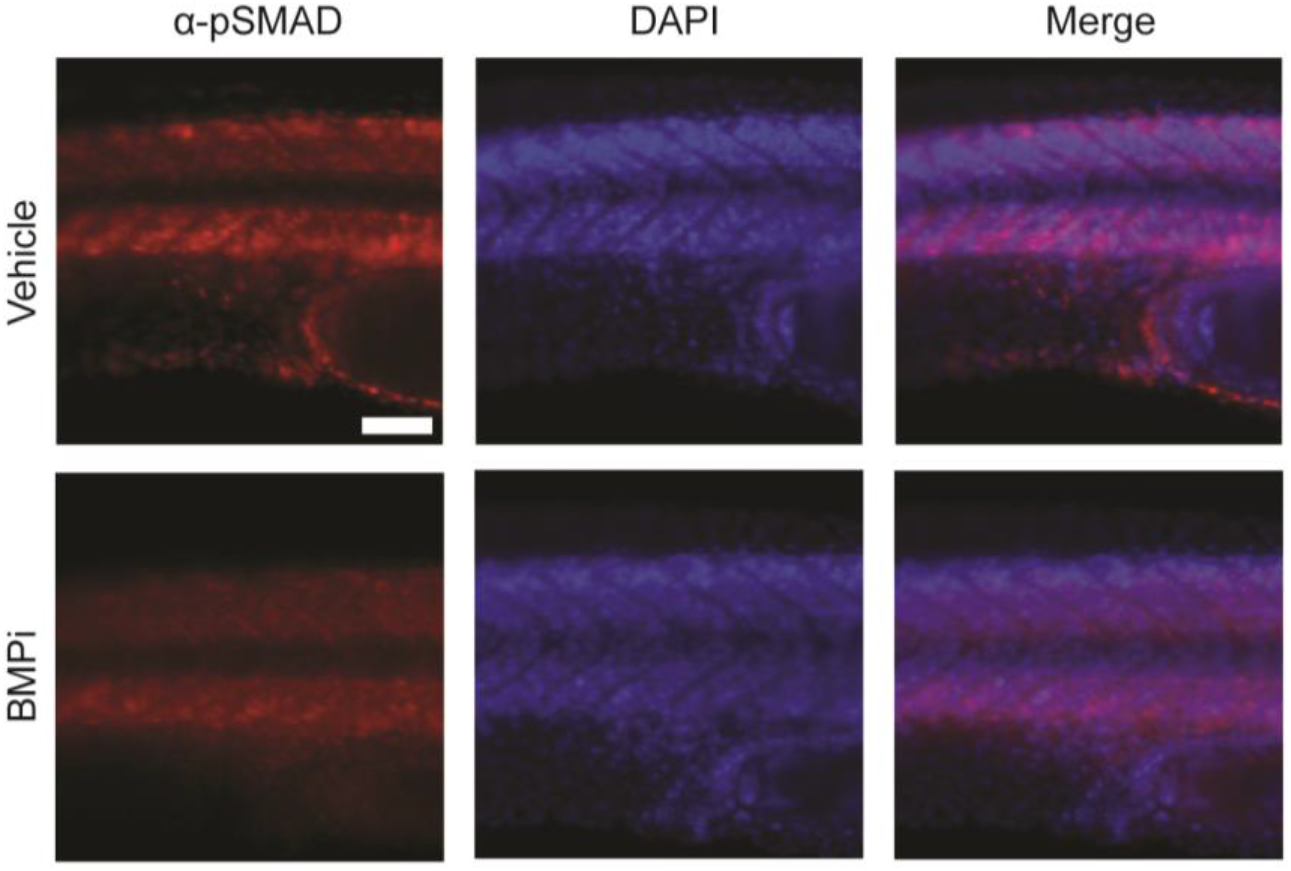
Treatment with the BMP inhibitor DMH1 reduces phospho-SMAD1/5/8 staining in embryos. Top, vehicle-treated animals and, bottom, BMPi-treated animals. Photomicrographs for pSMAD-1/5/8-stained embryos were taken at the same exposure settings. Scale bar = 50 µm.

**Figure S4 (Related to Figure 5).**
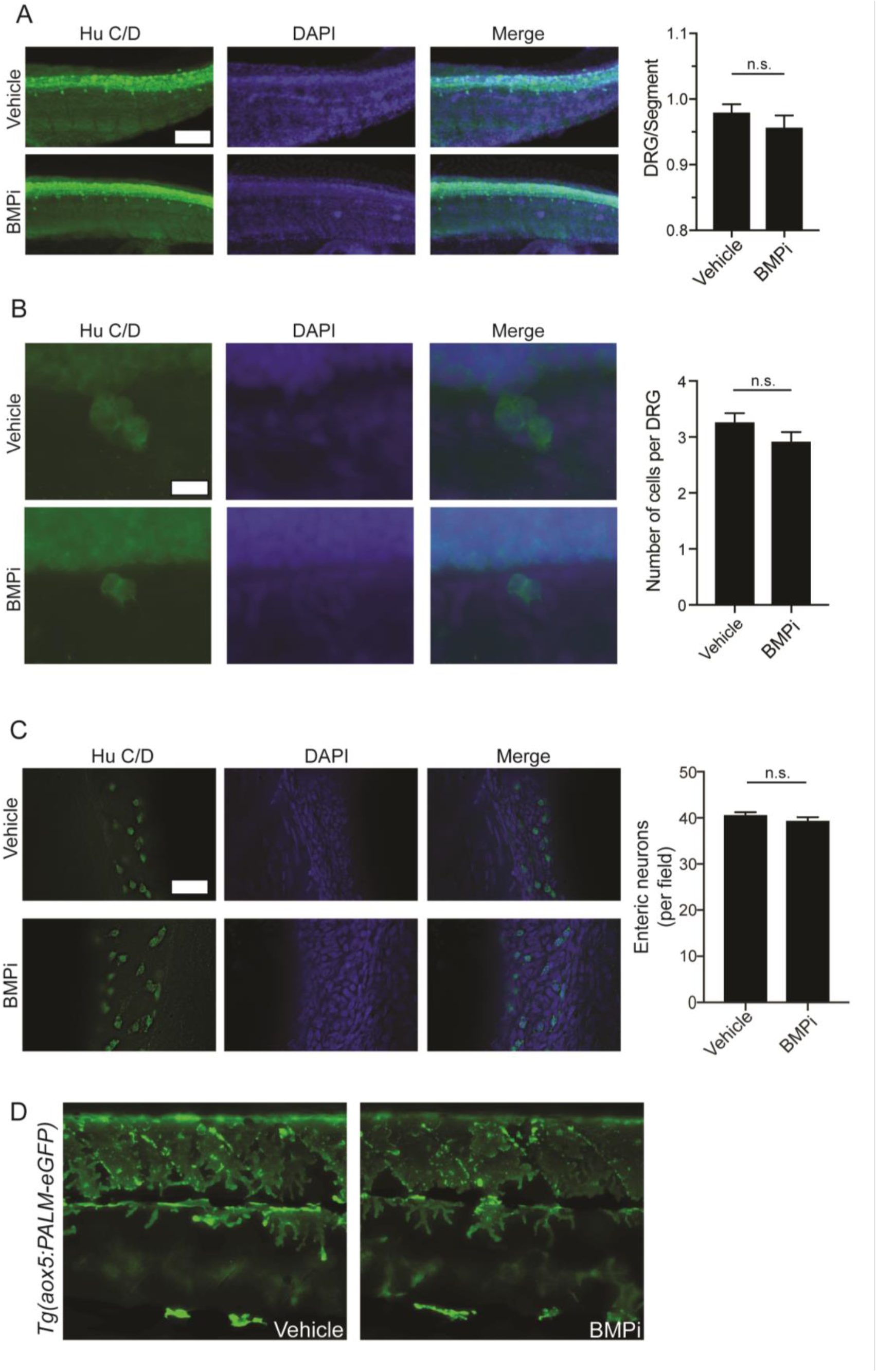
Some neural crest populations are unaffected by BMP inhibition. (A) Hu C/D staining for dorsal root ganglion structures showed no significant change in the number of dorsal root ganglia developing per segment, n = 5 per group. Scale bar = 50 µm. (B) Hu C/D staining for individual DRG’s showed no significant change in the number of cells populating each individual DRG; n = 29 per group. Scale bar = 10 µm. (C) Hu C/D staining for enteric neurons showed no significant change in number of enteric neurons per field in developing gastrointestinal tract; n = 4 per group. Scale bar = 50 µm. (D) Qualitative evaluation of xanthophore development using *Tg(aox5:PALM-eGFP)* embryos treated with vehicle or BMPi showed no apparent change in density or localization of xanthophores between vehicle- and BMPi-treated embryos, supported by no change in *aox5* expression as shown in Figure 5A. Error bars represent mean +/-SEM. P-values were calculated using Student’s t-test, n.s., not significant.

**Figure S5 (Related to Figure 6).**
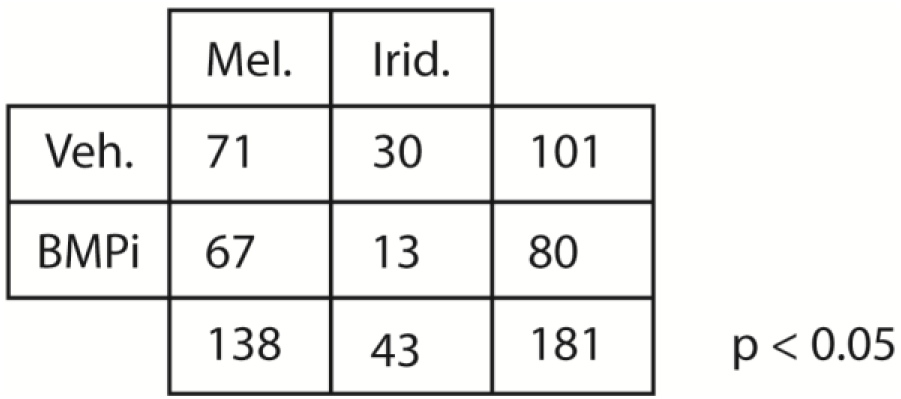
Quantification of iridophore and melanocyte numbers from lineage tracing. Number of iridophores and melanocytes identified by lineage tracing under each condition. P-value was calculated using Fisher’s exact test, P < 0.05.

